# Wheat mutants lacking Starch Synthase 1 have altered starch composition and cell wall content

**DOI:** 10.64898/2026.02.24.707672

**Authors:** Kay Trafford, Brendan Fahy, Oscar Gonzalez, Marianna Pasquariello, Jennifer H. Ahn-Jarvis, Judith Mayne, Ondrej Kosik, Alison Lovegrove, Frederick J. Warren, Brittany A. Hazard

## Abstract

Starch Synthase 1 (SS1) participates in the synthesis of amylopectin. To determine its role in hexaploid bread wheat, we selected and combined TILLING mutations for each homoeologue to create two independent SS1*-*deficient lines. The lines, which have different combinations of *ss1* mutations, both lacked SS1 protein. Both lines exhibited mild but significant changes to starch phenotype, including fewer short amylopectin chains, a slight increase in B-type starch granules and a modest increase in amylose content. Lack of SS1 also led to changes in the thermal properties of starch measured by differential scanning calorimetry including reduced enthalpy, and increased gelatinization temperature. Despite the changes to starch properties, the starch contents of the mutant lines compared to wild types were within the normal range, as were grain weight and protein content. However, the concentration of total- and water-extractable arabinoxylan, and MLG, were increased in white flour compared to wild-type controls.

**Highlight:** Lack of SS1 led to changes in starch molecular structure and thermal behavior. Starch content and grain weight were normal but cell wall polysaccharides in white flour were increased.

## Introduction

Wheat (*Triticum aestivum* L.) is one of the most widely-consumed global crops (Erenstein *et al*., 2022). As with other cereals, the major component of wheat grain is starch (approximately 80 % dry weight (DWT)) (Hazard *et al*., 2020). Whilst wheat’s distinctive storage protein, gluten, plays a critical role enabling the formation of dough used in bread and various other foods, properties of starch (such as granule size distribution; van Rooyen et al., 2023) have also been shown to influence the quality and acceptability of baked products.

Starch is composed of two glucose polymers, amylose (20 – 30 % DWT) and amylopectin (70 – 80 % DWT), which differ in molecular size, chain length, and branching patterns. Amylose is largely linear, made up of α-1,4-linked glucose units. In contrast, amylopectin is much larger and highly branched with α-1,6-linked branch points, with around 4–5 % of its glucose residues forming branch points. The branch points of amylopectin are clustered, and this facilitates crystallization, which in turn enables the formation of insoluble starch granules. Normal cluster formation and crystallization require amylopectin chains of different lengths. These are produced by the actions of three different isoforms of Starch Synthase (SS) named SS1, SS2 and SS3. Although their roles partially overlap (e.g. SS1 and SS3a in rice; Fujita et al., 2011), studies of mutants lacking individual SS isoforms have shown that SS1 is mainly responsible for synthesising the shortest chains, SS2 for the medium length chains and SS3 for the longest chains of amylopectin (Botticella *et al*., 2018; Fahy *et al*., 2022; McMaugh *et al*., 2014).

Most plant mutants lacking SS1 have fewer short amylopectin chains. In *Arabidopsis* (Delvallé et al., 2005) and rice (*Oryza sativa*) (Fujita *et al*., 2006), loss of SS1 resulted in amylopectin with proportionally less of the shortest glucan chains. In rice, the endosperm starch also had an elevated gelatinization temperature (Fujita *et al*., 2006). In contrast, Kossmann *et al*. (1999) reported no major changes in amylopectin structure in the tubers of transgenic potatoes with a near-complete reduction in SS1 activity. The leaf starch content of *Arabidopsis* mutants lacking SS1 (Delvallé et al., 2005) was reduced whereas in the ss1 mutant rice (Fujita *et al*., 2006), grain size was normal.

To date, only one study has examined the impact of reducing SS1 in wheat. RNA interference (RNAi) was used to suppress *SS1* gene expression specifically in the endosperm (McMaugh *et al*., 2014). This resulted in a reduction of SS1 protein in both the soluble phase and within starch granules, leading to alterations in chain-length distribution that were consistent with the findings in *Arabidopsis* leaves and rice endosperm. Reduction of SS1 in wheat endosperm was also shown to alter starch functional properties including gelatinization temperature, swelling power and starch viscosity.

To determine whether loss-of-function ss1 mutants display comparable properties to the previously characterized SS1 RNAi-suppressed wheat lines, and to provide valuable genetic resources for future investigation of the influence of SS1 on breadmaking quality, and for wheat breeding, we used a chemically induced TILLING (Targeting Induced Local Lesions IN Genomes) mutant population to generate SS1-deficient bread wheat cv. Cadenza. The production of knockout mutants in wheat is challenging due to the masking effects of functional gene copies in all three homoeologous genomes (A, B and D) (Borrill *et al*., 2015). However, advances in wheat genomic resources, particularly exome capture and re-sequencing of wheat TILLING populations, have enabled the identification of induced mutations in different homoeologues. Stacking these mutations by multiple genetic crosses facilitates the functional dissection of individual proteins (Krasileva *et al*., 2017). The impact of background mutations on phenotype can be mitigated by including multiple independent mutant lines and by comparing with wild-type controls. Here, we present the development and characterization of two SS1-deficient mutant wheat lines and their respective wild-type controls.

## Materials and methods

### Plant materials

For each *SS1* homoeologue (A, B and D), TILLING mutants carrying premature stop mutations (Table S1) were identified in the hexaploid wheat variety Cadenza using the in-silico Wheat TILLING database (www.wheat-tilling.com) and ordered from the JIC Germplasm Resources Unit (https://www.seedstor.ac.uk). Information on the TILLING mutants has since been incorporated into the Ensembl Plants variant database (https://plants.ensembl.org/Triticum_aestivum). The mutant plants were crossed, and two triple mutant lines, SS1-16 and SS1-17 (and their wild-type controls) were selected from the F_2_ progeny of triple heterozygous F_1_ plants, as described in Supplementary Fig. S1. Selections were made using KASP v4.0 genotyping (LGC Biosearch Technologies, Teddington, UK; www.biosearchtech.com) with homoeologue-specific primers (Table S2).

Plants were bulked up initially in a glasshouse and then grown in 1-m^2^ field plots harvested in 2023 (Bawburgh, Norwich, UK 52.627900, 1.179438). Six replicates per line were grown in a randomised complete block design. Further bulking was achieved in triplicate 12-m^2^ plots harvested in 2024 (Morley, Wymondham, UK 52.559430, 1.031256). Standard farm pesticide and fertiliser applications were made to reproduce commercial practise.

### Analysis of starch-granule-bound proteins

Samples of starch (25 mg) extracted from mature grains were suspended in 0.5 ml NuPage LDS sample buffer (Invitrogen, Paisley, UK; www.invitrogen.com), incubated at 95°C for 10 min with shaking at 750 rpm, allowed to cool on ice and centrifuged at 13,000 *g* for 5 min at 4 °C. The supernatants were removed and loaded onto 7 % gels (Tris-Acetate NuPage Mini gels, 1-mm thick, 15 well). Gels were run with SDS running buffer and stained (using SimplyBlue SafeStain) according to the manufacturer’s instructions.

### Analysis of soluble and granule bound proteins

Samples of developing wheat grains (average weight ∼40 mg, harvested approximately 10 days after anthesis) were extracted in ice-cold 100 mM MOPS, 1 mM DTT (1 µl/mg grain weight). Soluble and insoluble fractions were separated by centrifugation at 10,000 *g*, 4 °C for 10 min. Extraction buffer was added to the insoluble fraction to give the same final volume as the equivalent soluble fraction, loading buffer was added and the samples were heated at 70 °C (soluble) or 90 °C (insoluble) for 10 min. After cooling, equal volumes of soluble and insoluble fractions from each extract were loaded onto a 7 % 15-well NuPAGE®Tris acetate LDS precast polyacrylamide gel which was run according to the manufacturer’s instructions.

For Western blots, proteins were transferred onto nitrocellulose membranes by electroblotting using an Xcell II Blot Module (Thermo Fisher Scientific, Altrincham, UK; **Error! Hyperlink reference not valid.** following the manufacturer’s instructions. Blots were washed two times, each for 10 min, with 20 ml 150 mM NaCI and 10 mM Tris/HCI, pH 7.4, 0.1 % Tween 20 (TBST). After blocking for 5 h at room temperature in 20 ml TBST, 1 % BSA, 3 % skimmed milk powder, blots were probed overnight with primary antisera (maize anti-SSI; Mu et al., 1994) diluted 1:3000 by addition to the blocking solution. After washing three times, each for 10 min, with 20 ml TBST, the blots were incubated for 1 h with secondary antibody (goat anti-rabbit IgG alkaline phosphatase conjugate, Bio-Rad) diluted 1:5,000. Bands were detected using the 5-bromo-4-chloro-3-indolyl phosphate/nitroblue tetrazolium reagent system (Bio-Rad Laboratories Ltd., Watford, UK; bio-rad.com/en-uk).

### Grain morphology and composition

Morphometric parameters (grain width, length, area, and thousand grain weight) were measured for 400–600 grains per field plot (2023) using a MARVIN grain analyser (MARVITECH LTD, Stockport, UK; www.marvitech.de). Grain composition was determined for the same samples using a DA 7250 Perten Benchtop NIR Analyzer (PerkinElmer (UK) Limited, Beaconsfield, UK; www.perkinelmer.com). The NIR Analyser predicts composition from spectral measurements of whole intact grains. It uses wheat-specific calibration curves provided by the manufacturer that were produced from reference measurements of grain components compared with NIR spectra (Table S3B).

Mature grains were sectioned into 2-µm slices using a glass knife and microtome. Sections were mounted in water on glass slides, air-dried, and stained with 3% Lugol’s iodine solution (Sigma-Aldrich, Gillingham, UK; www.sigmaaldrich.com) prior to imaging. Microscopy was performed using either a 702 DM6000 (Leica) or an AxioObserver Z1 (Zeiss) instrument.

### Preparation of wholemeal flour and analysis of composition

Wholemeal flour was prepared by milling sub samples of grains (∼40-ml of grains by volume, each from a different 2023 field plot) using an UDY Cyclone Mill (UDY Corporation, Fort Collins, USA; www.udyone.com) fitted with a 0.5 mm screen.

Starch content was measured using thermostable alpha amylase and amyloglucosidase from the Total Starch HK (Hexokinase) Assay Kit (Megazyme Ltd., Bray, Ireland; www.megazyme.com) using a scaled-down protocol, as follows. Four replicate samples of each wholemeal flour (5-10 mg) were weighed into screw-capped, 1.5-ml tubes. Ethanol (20 µl of 80 % ethanol in water) was added and mixed with the flour for 30 s. To determine the free glucose plus starch content, three replicate tubes were digested with alpha amylase and amyloglucosidase. To each tube, 460 µl of 100 mM sodium acetate (pH 5.0) containing 90 units of thermostable alpha amylase was added and the tubes were incubated at 99 °C for 10 min in a thermomixer at 1,000 rpm. After cooling to 50 °C, 20 µl (66 U) of amyloglucosidase was added and the tubes were incubated at 50 °C for 35 min at 1,000 rpm. The fourth replicate flour sample was used to determine the free glucose content. This incubation was as above except that no amylase was added and 20 µl 100 mM sodium acetate (pH 5.0) was added in place of the amyloglucosidase. All digests were centrifuged for 10 min at 14,000 *g* and then 20 µl of the supernatant was diluted 10-fold prior to assaying 5 µl for glucose. Assays were performed in triplicate in 96-well plates in a total assay volume of 200-µl. Each assay contained 0.28 U hexokinase, 50 mM Hepes/NaOH (pH 7.5), 1 mM MgCl_2_, 1 mM ATP, 1 mM NAD and reactions were started by addition of 0.5 U glucose 6-phosphate dehydrogenase (NAD-specific from Leuconostoc) (Sigma-Aldrich). The initial and final optical densities at 340 nm were monitored using a plate reader (using the inbuilt conversion function to give absorbance readings in 1-cm pathlength-equivalents).

Carbon and nitrogen contents were measured using the Dumas combustion method. Sample of wholemeal flour (0.5 mg to 4 mg) were weighed using a Sartorius CUBIS II ultra micro balance (Sartorius, Göttingen, Germany) and analysed with a Leco TruSpec Micro CHN nitrogen analyser (Leco, Kirchheim, Germany; www.lecomet.eu). A conversion factor of 5.7 was used to calculate crude protein content.

### Preparation of white flour and measurement of cell wall polysaccharides

White flour was prepared by milling grains conditioned to 15.5 % moisture in a Metefém mini mill (METEFÉM Kft., Budapest, Hungary) (3-5 g, 2023 grains), followed by sieving to obtain the <150 µm white flour fraction or by the Allied Technical Centre (ATC, Maidenhead, UK; www.atcentre.co.uk) using a laboratory Buhler mill (Buhlergroup, London, UK;www.buhlergroup.com) (∼ 5 kg, 2024 grains). Total arabinoxylan (TO-AX) and water-extractable arabinoxylan (WE-AX) contents of white flour were determined by analysis of monosaccharides, and the relative amounts of enzyme-accessible AX and mixed-linkage glucans contents were measured by enzyme fingerprinting.

The monosaccharide content of white flour samples was measured using high-performance anion exchange chromatography (HPAEC; Dionex ICS-6000+; Thermo Fisher Scientific, Cambridge, UK; **Error! Hyperlink reference not valid.** For total monosaccharides, sugars were released from 200 µg aliquots of white flour by acidic hydrolysis in 2 M trifluoroacetic acid, heated to 120 °C for 60 min and then dried by centrifugation under vacuum, followed by washing with deionized water and drying. For water-extractable monosaccharides, 5 to 10 mg were first resuspended in Milli-Q water at a concentration of 5 mg/ml, mixed and left on a horizontal roller for 30 min. After centrifugation at 2,500 g for 10 min, 500 µL (equivalent to 2,500 µg of white flour) was transferred into a fresh screw-capped vial, dried and then hydrolysed with acid as above for total monosaccharides. Calibration curves were generated by subjecting authentic commercially available sugars, at a range of concentrations, (Sigma-Aldrich) to acid hydrolysis following the same protocol as for flour samples. Dried samples and standards were dissolved in 500 µl of 9 µM 2-deoxy-galactose in purified water (internal standard), and filtered through 0.45 µm PVDF disposable filters (Whatman, Maidstone, UK; cytivalifesciences.com/whatman) and chromatographed on a CarboPac PA20 analytical column (3 x 150 mm) with CarboPac PA20 guard column (3 x 30 mm) at 30 °C using an ICS-6000+ HPAEC equipped with an eluent generator with an EGC 500 KOH cartridge (Thermo Fisher Scientific). Twenty-five μl of diluted sample was injected onto the column. The total run time was 25.5 min, and the flow rate was 0.5 ml/min. The run conditions were: 0-16.5 min, isocratic at 3 mM KOH; 17–20 min, 100 mM KOH; 20.5–25.5 min, 3 mM KOH. Chromatograms were analysed and data calculated using Chromeleon 7.3 software (Thermo Fisher Scientific). Total-A+X and WE-A+X content were each calculated as the sum of arabinose and xylose.

Enzyme fingerprinting to measure enzyme-accessible AX and MLG was as described by Kosik *et al*. (2017) with minor modifications. Briefly, after boiling and washing the white flours with ethanol, these were digested with 2 µL of recombinant endo-1,4-xylanase 11A (NpXyn11A; Prozomix, Haltwhistle, UK; prozomix.com) and 1 µL of lichenase (CtGH26; Prozomix) in 0.1 M sodium acetate buffer pH 5.5 at 40 °C with shaking overnight. Enzymatic digestion was terminated by boiling the sample for 30 min. Samples were centrifuged at 13,400 *g* for 5 min before being filtered through 0.45 µm PVDF and diluted 1:20 with 7 µM melibiose in purified water (internal standard). The enzymatically-released arabinoxylan oligosaccharides (AXOS) and mixed-linkage glucan oligosaccharides (MLGOS) were separated using a CarboPac PA1 analytical column (2 × 250 mm) with CarboPac PA1 guard column (2 x 50 mm) (Thermo Fisher Scientific) at 30 °C with a flow rate of 0.25 ml/min, following the original method of Ordaz-Ortiz, Devaux and Saulnier (2005). The areas under the AXOS peaks were combined to determine total enzyme-accessible AX and areas under the G3 and G4 (the major MLG-derived oligosaccharides in white flour) MLGOS peaks to give mixed linkage glucan content (both expressed in arbitrary units). Samples were analysed in triplicate.

### Starch purification

Starch was purified from single mature grains as follows. Each grain was soaked in 300 µl distilled water at 42 °C for 1 h then cooled on ice. Grains were cut into small pieces with scissors and then ground to homogeneity in a 1.5-ml tube using a plastic pestle and an electric mini-drill. Two 150-µl sub samples of well-mixed homogenate were each layered on top of 1,350 µl ice-cold 80 % (w/v) CsCl in water in 2-ml tubes and centrifuged at 13,000 *g*, 4 °C for 5 min. The supernatant was carefully removed and discarded leaving a white starch pellet. The pellet was washed three times by resuspension in 300 µl water, centrifugation at 13,000 *g*, 4 °C for 5 min and removal of the supernatant. Starch pellets from the two 150-µl sub-samples were combined together before the final water wash. The final starch pellets were stored at -20 °C.

Starch was extracted from 25-50 mg white flour as for single grain extracts (except that the soaking at 42 °C step was omitted). When larger samples of starch were required, three replicate grains or white flour samples were extracted as above and then the starch pellets were combined before the final water wash. If dry starch was required, the final pellets were resuspended in a small volume of acetone, centrifuged and the acetone removed and discarded. The pellets were then left to dry completely in a fume cupboard overnight.

Starch was purified from 2.5-g samples of wholemeal flour by grinding in excess ice-cold water in a mortar then filtering through one layer of Miracloth (Merck, Haverhill, UK; www.merckgroup.com/uk-en) into a 50-ml tube. After washing through the filter cloth with additional water, the filtrate was centrifuged at 500 *g*, 4 °C for 5 min and then the supernatant was discarded, and the pellet was resuspended in 30 ml of 2 % (w/v) SDS. The centrifugation and supernatant-removal steps were repeated once more. The pellet was transferred to a 15-ml tube and washed (as above), twice with 10 ml of 2 % (w/v) SDS, and twice with 5 ml water, discarding the supernatants after each resuspension and centrifugation step. The starch was resuspended in 5 ml water and replicate sub-samples of 0.3 ml were each loaded on to 1.5 ml of 80 % (w/v) CsCl in water in a 2-ml tube and centrifuged at 13,000 *g* for 5 min. The supernatant, and any non-starch material, was removed from the starch pellet and discarded. After two washes with 300 µl water, the starch was resuspended in 300 µl of absolute ethanol, centrifuged at 13,000 *g* for 5 min, and the supernatant discarded. The purified starch was air-dried overnight in a fume hood.

### Starch-granule morphology and size distribution

The morphology of starch purified from wholemeal flour was examined using a Zeiss Gemini SEM300 scanning electron microscope (Zeiss, Cambourne, UK; www.zeiss.co.uk) by the JIC bioimaging staff. Prior to microscopy, a small amount of starch was added to 50 µl absolute ethanol and after mixing, 2-µl aliquots were added to a round coverslip attached to an SEM stub, allowed to dry and then sputter-coated to 8 nm thickness with gold.

Particle-size distribution was determined using a Multisizer 4e Coulter counter (Beckman Coulter, High Wycombe, UK; beckmancoulter.com) fitted with a 70-µm aperture tube. Purified starch was suspended in Isoton II electrolyte solution (Beckman Coulter). After filtration using a 70-µm cell strainer, a minimum of 100,000 particles per sample were measured. These data were used to produce relative volume vs. diameter plots. To calculate the mean diameters of the A- and B-type granules and the B-type granule volume percentage (defined as volume occupied by B-type granules as a percentage of the total volume of starch), a mixed bimodal distribution (log normal/normal) was fitted to the plots (Python script available at https://github.com/DavidSeungLab/Coulter-Counter-Data-Analysis).

### Analysis of starch molecular structure

The amylose contents of purified starches were measured using an iodine-binding assay based on that described by Knutson and Grove (1994). A 0.05% solution of starch in DMSO containing 6 mM iodine was prepared and mixed overnight at room temperature. Samples were diluted ten-fold with water and the absorbance at 600 nm was measured using a spectrophotometer. Amylose content was determined by comparison with potato amylose type III standards (Sigma-Aldrich). To allow for iodine binding by amylopectin, the apparent amylose values were corrected using the following equation: % amylose = (% apparent amylose-6.2)/ 93.8.

Debranched starch samples were prepared as follows. Purified starch (1 mg) was weighed into a screw-capped, 2-ml tube, suspended in 1 ml water and heated to 121 °C for 15 min. After cooling, 50 µl 50 mM sodium acetate buffer (pH 3.5) was added and 250 µl of the mixture was transferred to a screw-capped 1.5-ml tube together with 5 µl (2.5 U) of isoamylase (E-ISAMYHP; Megazyme) and incubated at 37 °C for 18 h. The reaction was stopped by heating to 99 °C for 2 min. A 50-µl sample was diluted 10-fold and filtered by adding to 450 µl water in a 0.22 µm Millipore spin filter (UFC 30GV00; Sigma-Aldrich) and centrifuging at 12,000 *g* for 5 min.

The molecular-weight distributions of debranched starches were measured using size-exclusion chromatography on PSS 300 and 3000 columns in series, with a guard column (Agilent-Technologies, Mainz, Germany; www.agilent.com). The columns were connected to an HPLC (Walters Alliance; Waters Ltd., Wilmslow, UK; www.waters.com) equipped with a refractive index detector, autosampler (40°C), and column heater (80°C). The HPLC was run isocratically at a flow rate of 0.6 ml/min using 0.5 % (w/w) LiBr/DMSO as the eluent.

Samples were injected at a concentration of 1 mg/ml. Molecular mass was determined by comparison with unbranched pullulan standards (Standard P-82) (Shodex, Munich, Germany; www.shodex.de).

The molecular-weight distributions of debranched starches were also measured using anion-exchange chromatography on a CarboPac™ PA-100 analytical and guard column (Thermo Fisher Scientific). Short glucan chains were separated and detected using an HPLC equipped with a pulsed amperometric detector (Dionex ICS-3000 HPLC; Thermo Fisher Scientific). The columns were equilibrated with buffer A (107.5 mM NaOH, 75 mM sodium acetate) and eluted with a gradient from 100 % buffer A to 100 % buffer B (130 mM NaOH, 300 mM sodium acetate) over 45 ml at flow rate of 1 ml/min. The debranched starch peaks were identified by comparison with glucan standards of 1-7 glucose units.

### Analysis of the functional properties of starch and wholemeal flour

The swelling power of purified starches in 1 % SDS and in water was measured as follows: 10 mg of starch was added to a pre-weighed 2-ml round-bottomed screw-capped tube. After addition of 1 ml of water (or 1 % SDS), the tube was incubated at 80 °C for 30 min with shaking at 750 rpm and then centrifuged at 1,500 *g* for 5 min. The supernatant was carefully removed and discarded and the tube plus starch pellet was weighed. Swelling power was calculated by dividing the weight of the starch after swelling by the weight of the dry starch.

The loss of birefringence of starch was examined using a light microscope fitted with a heated stage, λ-plate, polarizing filters and a 10x objective. Starches were SS1-17 mutant or wild-type controls, purified from white flour from grains harvested in 2024. Each starch sample was suspended in dilute Lugol’s solution. The slides were heated at 5 °C/min from 25 to 90 °C and images were captured at specific temperatures during heating.

Differential scanning calorimetry was performed using a DSC2500 (TA Instruments, Elstree, UK; www.tainstruments.com). Starch (1 mg) was accurately weighed into an aluminium pan (Tzero, TA Instruments) then 10 µl of degassed water was added and the pan lid hermetically sealed. The furnace was purged with dry nitrogen gas at a flow rate of 50 ml/min. The measurement procedure started with temperature equilibration at 10 °C for 10 min, followed by a temperature ramp up to 85 °C at the rate of 2 °C per min. DSC parameters were analysed using TA Instruments’ software package, Trios.

Digestibility of wholemeal flour by pancreatic alpha-amylase (Sigma-Aldrich) was measured in vitro. Wholemeal flour (∼12.5 mg) was accurately weighed into 1.5-ml round-bottomed, screw-capped tubes and Phosphate Buffered Saline (PBS; Merck) added to achieve 5 mg/ml starch. A 3-mm diameter glass bead was added, and tubes were incubated at 80 °C in a pre-heated 24-well thermomixer for 20 min at 1000 rpm. The gelatinized starch samples were cooled to room temperature and either assayed immediately (cooked) or incubated at 4 °C overnight (cooked and cooled) before assay. Subsamples of starch were transferred to the wells of a 96-well PCR plate for digestion. Up to eight flours were digested per 96-well plate, one per row. In each row, replicate 54-µl subsamples of gelatinized flour were added to wells 1-11 and 200 µl of a 10x stock solution of 2 U/ml of α-amylase (19.5 µmol maltose-equivalents/min/ml) was added to well 12. After preheating to 37 °C on a thermomixer for 10 min at 350 rpm, 60 μl of 300 mM Na_2_CO_3_ stop solution was added to the 0-min control samples followed by 6 μl of 10x amylase stock solution. The remaining samples were digested for different lengths of time by adding alpha amylase, mixing thoroughly with the pipette tip, then incubating for an appropriate time at 37 °C and 350 rpm before adding stop solution. The stopped digests were covered with flat-topped strip seals. Plates were centrifuged at 850 *g* for 10 min at 20 °C prior to assaying the supernatants for reducing sugars.

To assay reducing sugar content, 15-µl subsamples of each stopped digest was added to the corresponding well in an optical microplate and diluted by addition of 5 µl PBS. Eight maltose standards (0-60 nmol in 20 µl) were added to every plate (in column 12).

Immediately before use, a stock assay mix (Lever, 1975) was prepared as follows: 100 mg of PAHBAH (*p*-hydroxybenzoic acid hydrazide) was dissolved in 1.95 ml 0.5 M HCl, then 0.5 M NaOH was added to give a final volume of 20 ml. After adding 180 μl of stock assay mix to each well, the plate was covered with adhesive foil sealing film, incubated for 20 min in a thermomixer at 99 °C and 350 rpm, centrifuged at 1,500 *g* for 5 min at 20 °C (to cool and collect condensation) and then the optical density at 405 nm was determined immediately using a plate reader (VersaMax microplate reader; Molecular Devices, CA, USA; www.moleculardevices.com). The reducing sugar content (as maltose equivalents) was determined by reference to the maltose standards on the same plate.

The digestion-with-time curves for each flour were compared using logarithm of the slope (LOS) plots as described by Butterworth *et al*. (2012). For each pair of adjacent time points (t1 and t2), the difference in the corresponding reducing sugar concentrations (c1 and c2) was calculated. The LOS values were calculated as the natural logarithms of (C2− C1)/(t2− t1), (C3− C2)/ (t3− t2) etc. and were plotted against the mean time ((t2− t1)/2). To allow for the reducing sugar content of the zero min incubations, this time point was assumed to be 0.001 min. A linear curve was fitted to the LOS plot data and using this, the intercept (ln(C∞k)) and gradient (-k) were determined. From these values and the total incubation time (e.g. 80 min), C∞ and e^-ktx^-1 were calculated. Finally, for each flour digest, the AUC was calculated as (C∞ x total incubation time) + ((C∞ / -k) x (e^-ktx^-1)).

### Statistical analysis

The statistical significance was determined using Student’s t-tests. As well as giving P values, we used the asterisk rating system, as follows: P < 0.05*, P < 0.01**, and P < 0.001***.

## Results

### All three homoeologs of SS1 are expressed in wheat endosperm

Analysis of publicly available expression data (expVIP database) showed that all three homoeologs of SS1 are expressed in developing wheat endosperm, with the highest expression during the first half of endosperm development (6-14 days post anthesis) (Fig. S2). Between homoeologs, the relative expression levels differ approximately 2-fold in the order *SS1-7B*>*SS1-7D*>*SS1-7A*.

Analysis of natural variation in *SS1* sequences amongst elite varieties and landraces (https://plants.ensembl.org/Triticum_aestivum/) showed that there is one missense variant (Asn218Lys) at the end of exon 2 of *SS1-7A* (TraesCS7A02G120300) that is predicted to inactivate the protein (SIFT=0 i.e. deleterious). The variant SNP (G instead of C) results in a change of the encoded amino acid from asparagine (Asn) to lysine (Lys, codon AAG). The mutation is in the N-terminal starch synthase catalytic domain (a domain largely specific to SS enzymes) and is close to the active site cleft (AlphaFold predicted model, Ensembl Plants). This natural missense variant of *SS1-7A* may therefore affect protein content, enzyme activity and/or binding to starch.

The variant present in *SS1-7A* in the wheat reference variety, Chinese Spring, encodes asparagine (codon AAC). The *SSI-7B* and *SS1-7D* homoeologous genes of Chinese Spring also both possess Asn in the corresponding position. Comparison of the corresponding amino acid in other SS1 proteins (https://plants.ensembl.org) showed that most cultivated and wild-type Triticeae species, and related cereal species, (eg. *Hordeum vulgare*, *Brachypodium distachyon*, *Secale cereale*, *Aegilops tauschii*) possess the Asn variant. This suggests that the ancestral, presumably wild type, amino acid is asparagine and that the lysine variant is a more recent, possibly deleterious mutation. The variety Cadenza has the mutant lysine variant i.e. *SS1-7A* in Cadenza is possibly inactive.

### *ss1* TILLING mutants lack SS1 proteins

To create bread wheat mutant lines entirely lacking SS1 activity, nonsense mutations (with premature stop codons) were chosen from the available TILLING mutants in the variety Cadenza for each *SS1* homoeolog. Two lines were created, SS1-16 and SS1-17, that shared the same *ss1-7A* and *ss1-7D* mutant alleles but differed in their *ss1-7B* alleles (Fig. 1A). The *ss1*-*7B* mutation used in line SS1-16, and the *ss1*-*7D* mutation used in both lines, were positioned close to the C-terminal ends of the corresponding proteins, raising the possibility that slightly truncated (1-2 % reduced mass) SS1 proteins, retaining some activity, could be produced.

**Fig. 1.**
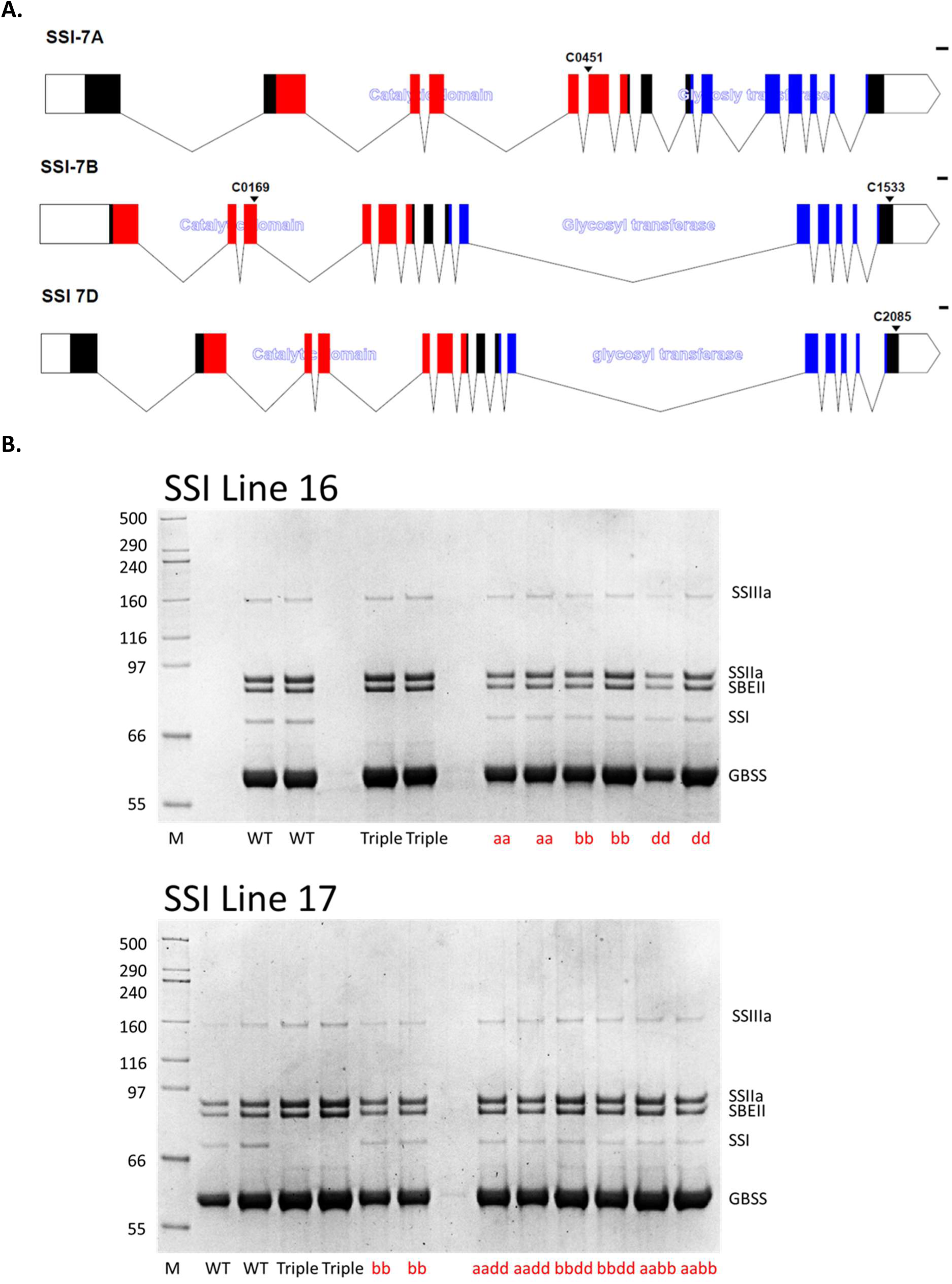
Starch Synthase 1 gene structures and gels of starch granule-bound proteins. **A.** The gene structures of the *SS1* homoeologs (as annotated in Ensembl Plants, release 57) were drawn using WormWeb exon-intron graphic maker (WormWeb.org). Unfilled bars represent UTRs. The catalytic (red) and glycosyl transferase (blue) domains, and the positions of the nonsense TILLING mutations are indicated. The scale bar is 100 bp. **B.** Starch granule-bound proteins were extracted from wild type and mutant (SS1-16 and SS1-17) grains and separated by SDS-PAGE. The molecular masses (kDa) and identities of the proteins are indicated. Note the absence of the SS1 protein in the triple mutants (Triple) and its presence in the wild type (WT), single (eg aa) and double mutants (eg aabb).

**Fig. 2.**
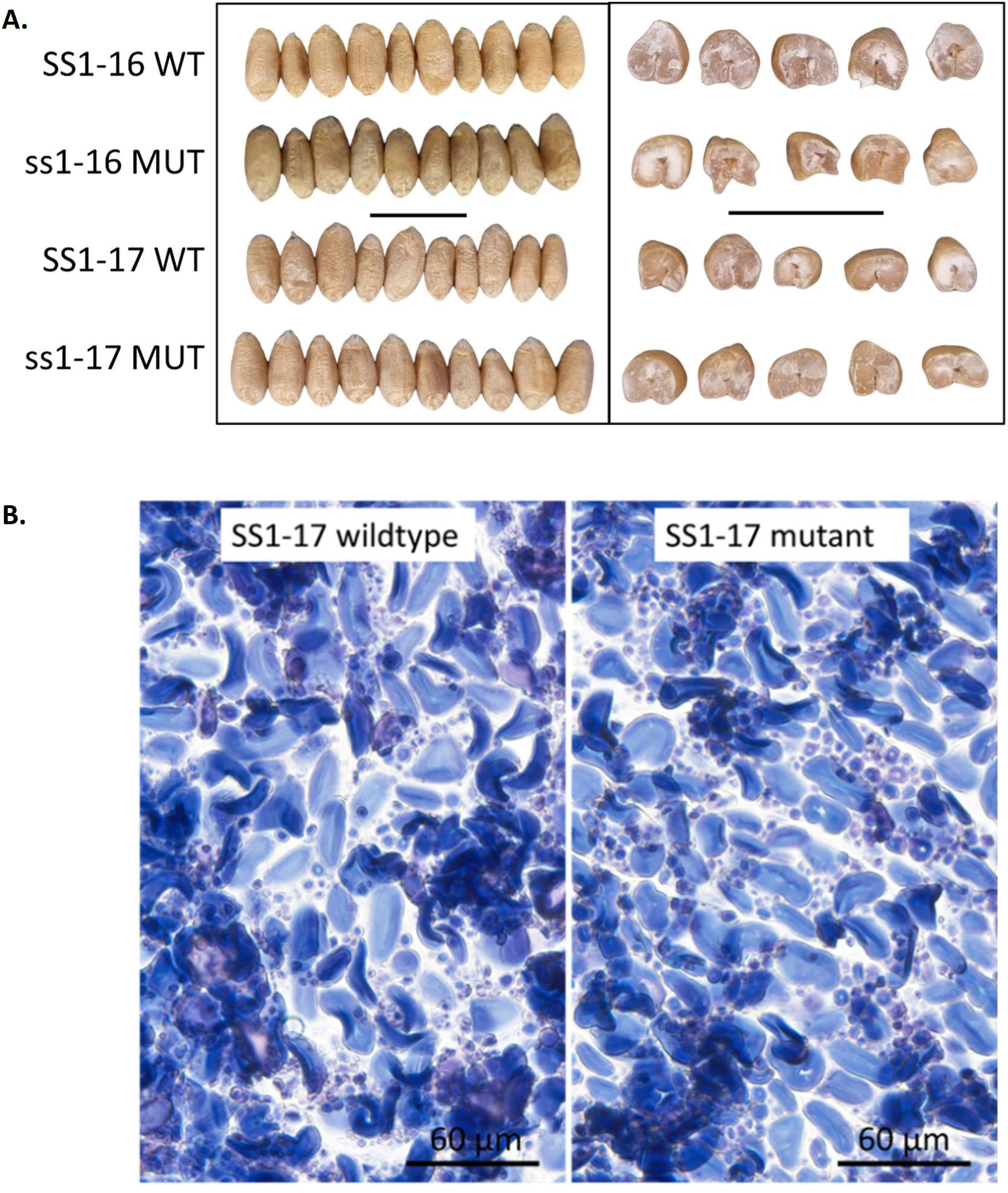
The appearance of mature grains and endosperm. **A.** Photographs of typical SS1 grains and grain segments from field-grown (2023) mutant (MUT) and wild-type (WT) plants. Scale bars are 1 cm. **B.** Light microscopy of endosperm sections from mature grains of line ss1-17 WT and MUT. Sections were stained with 3% Lugol’s iodine solution (Sigma-Aldrich, Gillingham, UK) prior to imaging. Scale bar is 60 µm.

SS1 was shown previously to be almost entirely present in the granule-bound fraction in developing maize endosperm (Mu-Forster et al., 1996). Since few proteins in the endosperm are starch-granule-bound like SS1, examination of the types of granule-bound protein using SDS-PAGE provides a means to assess the impact of both the nonsense TILLING mutations, and the natural missense variant of SS1-7A (Asn218Lys), in Cadenza on SS1 protein content and its ability to bind to starch. We examined the proteins embedded in starch granules isolated from mature grains using SDS-PAGE (Fig. 1B) and we compared the amounts of SS1 proteins in soluble and insoluble fractions of developing grains using Western Blots (Fig. S3).

In wheat starch, SS1 was identified as a 75 kDa band (Yan, Fairclough and Bhave, 2000). A band of this size was observed in all double mutant, single mutant and wild-type starches using SDS-PAGE but not in the triple ss1 mutant (aabbdd; Fig. 1B). No novel truncated SS1 proteins were observed in any of the mutants. Our Western blots showed that SS1 was detectable in the granule-bound fraction of developing wheat grains but not in the same proportion of soluble fraction (Fig. S3). Taken together, these results suggest that the three *SS1* homoeologues in Cadenza encode proteins of similar size and that these SS1 proteins are absent from both the ss1-16 and ss1-17 triple-null mutants. The AAbbdd double mutant lacks the *ss1-7A* TILLING mutation but possesses the natural missense variant Lys218. Since this double mutant lacks the SS1-7B and SS1-7D proteins, the SS1 protein present in the starch granule-bound fraction must be encoded by the *ss1-7A* missense variant. This indicates that the *ss1-7A* missense mutant possesses a full-length protein that is capable of binding to starch. We did not assess whether the missense variant protein possesses starch synthase activity. However, for simplicity, our further studies were restricted to the ss1 triple-null TILLING mutant grains. The triple nulls will be referred to as the ‘ss1 mutant’ and their wild-type controls as ‘ss1 wild type’.

### Mutants lacking SS1 have near-normal grain weight

No consistent visual differences between SS1 wild type and mutant grains (Fig. 1A) or starch granule (Fig. 1B) in size or shape were observed. Grain morphology was also determined using a Marvin seed analyser, and physical characteristics were determined using an NIR seed analyser (Table S3). For these studies, SS1 lines (SS1-16 and SS1-17) were grown in 2023 in randomized field plots consisting of six replicate plots for each mutant (aabbdd) and wild type control (AABBDD). One consistent difference only was found between the wild type and mutant ss1 grains: for both lines (16 and 17), the mutant grains had lower NIR-predicted hardness. Other differences between the wild type and mutant ss1 grains that were observed in one line but not in the other included: in mutant ss1-16, there was decreased predicted dry gluten and protein, and decreased grain weight whereas in mutant ss1-17 there was increased moisture content and decreased starch content.

### Mutant endosperm had increased concentrations of cell wall polysaccharides

Mature grains from the 2023 field plots were milled to produce wholemeal or white flour and its composition was determined biochemically (Table 1). There was no difference in starch content between ss1 wild type and mutant for either line. In mutant ss1-17, the protein content was increased, and the C/N ratio was decreased but in mutant ss1-16 these measurements were not significantly different.

**Table 1.** Grain composition. Six 1-m^2^ field plots each for SS1 wild type (SS1, AABBDD) and mutant (ss1, aabbdd) lines were grown in 2023. Each plot was a different sibling line. Samples of grains from each plot were milled to wholemeal or white flour. Six replicate wholemeal flours were assayed for starch (by enzymatic assay), and for protein content and C/N ratio (by Dumas combustion assay) and 3 replicate white flours were assayed for cell wall components. The starch content per grain was calculated using the grain weight data shown in Figure S3. Total arabinoxylan (TO-AX) and water-extractable arabinoxylan (WE-AX) contents were determined by analysis of monosaccharides and the relative amounts of Total AX and mixed linkage glucans (MLG, β-glucan) contents were measured by enzymatic fingerprinting. The mean values +/- SE are shown together with statistical analysis (Student’s t-tests) of the differences between wild type and mutant (p-values, ns = no significant difference).

| Sample | Starch<br>content<br>(% DWT) | Starch<br>content<br>(% FWT) | Starch<br>content<br>(mg/grain) | Protein<br>content<br>(% FWT) | C/N ratio | TO-AX<br>(mg/g<br>FWT) | WE-AX<br>(mg/g<br>FWT) | TO-AX<br>(arbitrary<br>units) | MLG<br>(arbitrary<br>units) |
| --- | --- | --- | --- | --- | --- | --- | --- | --- | --- |
|  | Wholemeal | Wholemeal | Wholemeal | Wholemeal | Wholemeal | White | White | White | White |
| SS1-16 | 68.2 ± 11 | 58.7 ± 09 | 30.8 ± 07 | 14.5 ± 01 | 15.9 ± 01 | 18.8 ± 06 | 4.55 ± 016 | 11.29 ± 032 | 1.744 ± 009 |
| ss1-16 | 68.0 ± 18 | 59.2 ± 16 | 28.4 ± 07 | 14.1 ± 01 | 16.2 ± 02 | 23.9 ± 06 | 6.71 ± 051 | 13.98 ± 070 | 2.155 ± 007 |
| t-test | 0.9235 ns | 0.7787 ns | 0.0544 ns | 0.3357 ns | 0.6175 ns | 0.0045 ** | 0.0161 * | 0.025 * | 0.0239 * |
| SS1-17 | 69.1 ± 14 | 61.5 ± 14 | 24.5 ± 22 | 13.3 ± 01 | 17.4 ± 01 | 18.8 ± 07 | 5.05 ± 009 | 11.65 ± 017 | 1.814 ± 003 |
| ss1-17 | 71.8 ± 30 | 62.1 ± 28 | 27.5 ± 13 | 14.1 ± 01 | 16.4 ± 01 | 23.1 ± 04 | 7.10 ± 024 | 14.25 ± 028 | 2.465 ± 009 |
| t-test | 0.4417 ns | 0.8482 ns | 0.2228 ns | 0.0291 * | 0.0193 * | 0.0057 ** | 0.0013 ** | 0.0014 ** | 0.0026 * |

For both ss1 lines, the NIR analysis of mature grains showed no significant difference in predicted Neutral Detergent Fibre content (NDF) (Table S3). NDF content is a measure of the total plant cell wall content. Most of the NDF in whole grains is in the testa and pericarp (husk) which, together with the embryo, is largely removed (as bran) when wheat grains are milled to white flour. Therefore, white flour consists mainly of wheat endosperm. Since the fibre content of white flour is a trait of interest, we assayed the contents of the two major endosperm cell wall polysaccharides: arabinoxylan and mixed-linkage glucan (MLG or β-glucan). Samples of white flour were prepared from three of the six replicate field plots (2023) and analysed for fibre content using two different methods. For both ss1 mutants, there were significant increases in the amounts of arabinoxylan (total and water-extractable) and β-glucan in white flour compared with their wild-type controls (Table 1).

### Mutants lacking SS1 have near-normal starch granule morphology

Wheat starch consists of two distinct granule types: larger, disc-shaped A-type granules (∼20 μm in diameter) and smaller, more abundant spherical B-type granules (∼7 μm) (Chia *et al*., 2020). Light microscopy showed that most of the starch granules in mutant wholemeal flour (2023) had normal morphology (Fig. 3A). However, in the ss1 mutants, there were a few abnormal A-granules that looked distorted in the bright-field image and had an unusual pattern of birefringence in the corresponding polarized-light image. These distorted A-granules were also larger than normal.

**Fig. 3.**
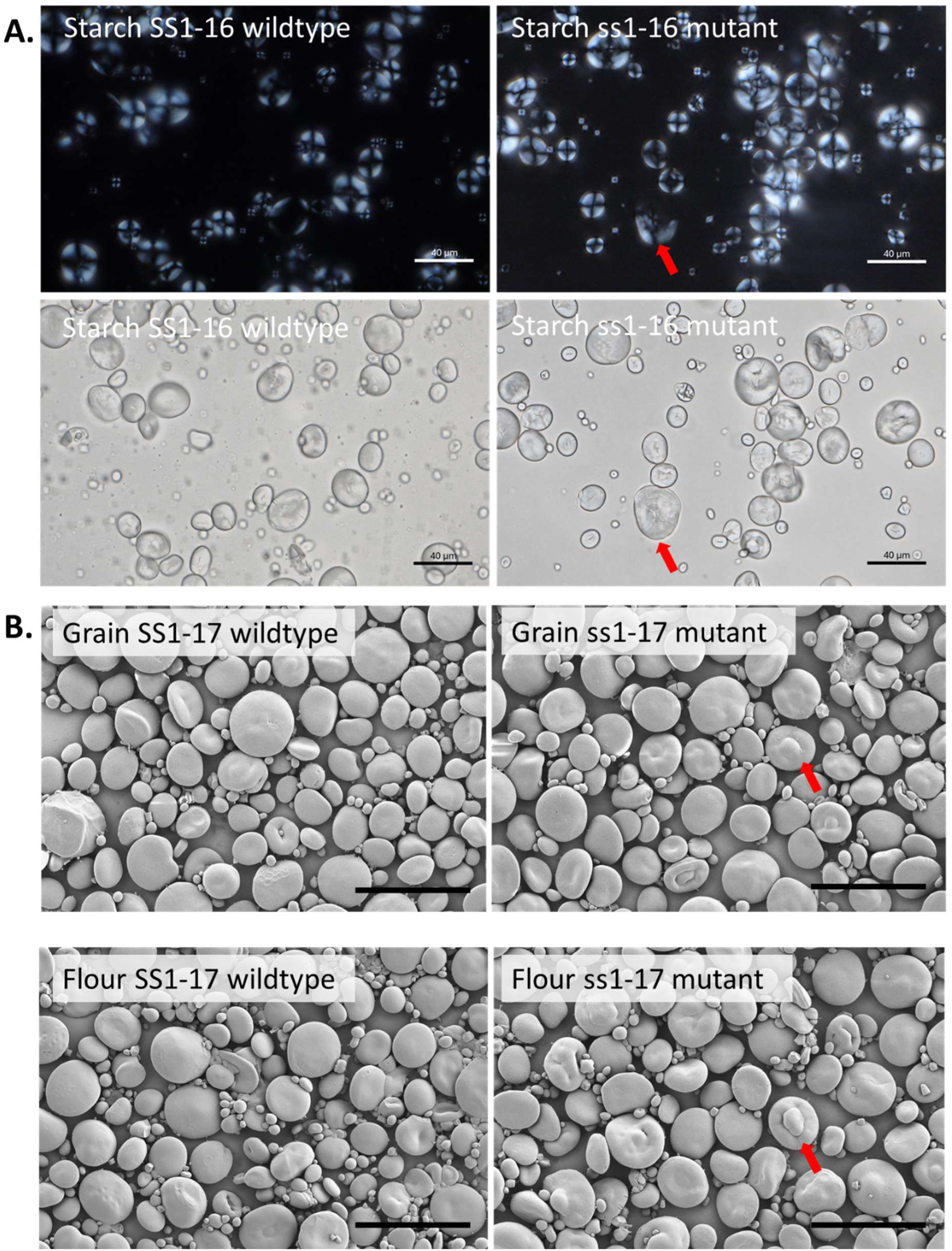
Starch granule morphology. SS1-16 and SS1-17 mutant and wild type plants were grown in field plots in 2023 and 2024. **A.** Wholemeal flours from 2023 grains were observed using light microscopy, with and without a polarizing filter. The results were similar for both lines. Only images for SS1-16 are shown. The arrows indicate a granule that has partially lost its birefringence. The scale bar is 40 µm. **B.** Starch was prepared from 2024 grains or white flour and imaged using an SEM microscope. Three separate grain and white flour samples prepared from each SS1-17 genotype were examined and the differences in the structure of the granules were consistent. The arrows indicate some of the granules with abnormal morphology. Abnormal granules were seen in all starches, but they occurred more frequently in starch from mutant grains that had been milled. The scale bar is 50 µm.

Starch granule-size distributions of starches purified from the wholemeal flour from field-grown grains (2023) were analysed using a Multisizer 4e Coulter counter (Fig. 4A and Table S4A). For both SS1 lines, the mutants had slightly more B-granules than their respective wild-type controls. There were no differences in A-granule diameter between the mutants and wild types. However, ss1-17 mutant had B-granules with a slightly greater diameter than wild type (Table S4A).

**Fig. 4.**
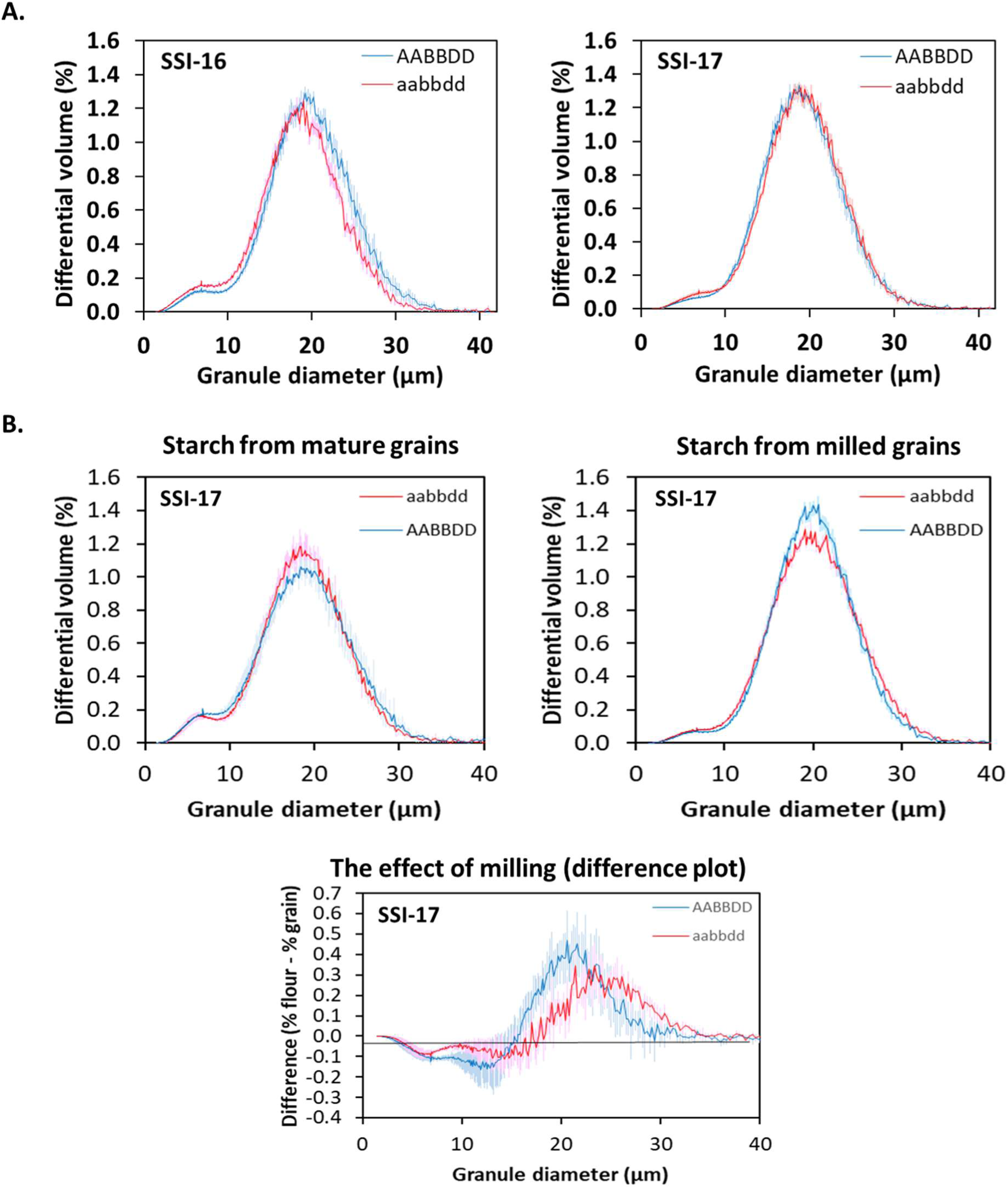
Starch granule-size distribution. Starch granule-size distributions were assessed using a Multisizer 4e Coulter counter. **A.** Starches were prepared from wholemeal flour from field-grown grains (2023). Values are means ± SE of six biological replicates, each from a different field plot. Values for A- and B-granule diameters and B-granule content are shown in Table S4. **B.** To assess the effect of milling on granule-size distribution, starch was extracted from the same batches of grains (field-grown, 2024) by two different methods: gentle extraction of rehydrated grains in aqueous solution or extraction after milling to white flour. Values are means ± SE of three biological replicates, each from a different field plot. The difference plot shows the effect of milling on granule-size distribution for wild type and ss1 mutant starches. The same starches were used for SEM (Fig 3B).

To quantitatively assess the effect of milling on granule-size distribution, starch from SS1-17 mutant and wild-type controls was prepared from field-grown grains (2024) by two different methods: gentle extraction of rehydrated grains in aqueous solution and extraction from grains that were first milled to white flour (Fig. 4B). Milling resulted in slightly different starch granule-size distributions. This could be seen most clearly using a difference plot. For both wild type and mutant, milling resulted in proportionally fewer B-type granules. Interestingly, in the mutant but not in the wild type, there was an increase in the size of the A-type granules after milling, with the presence of proportionally more granules of ∼25-35 µm diameter than in the gently extracted starches.

Scanning electron micrographs of starch granules purified from 2024 grains or white flour are shown in Figure 3B. Some A-granules from the ss1 mutant grains had bulbous, protruding central regions rather than the normal smooth, discoid shape. Such distorted granules were seen more frequently in white flour samples than in starches that were gently extracted from mature grains.

### Mutants lacking SS1 have reduced amylopectin content

To quantify the effect of SS1-deficiency on amylopectin content, the amylose contents of purified starches from glasshouse-grown plants were measured using an iodine-binding assay (Fig. 5A) and the amylopectin to amylose ratios of starches from the 2023 field plots were measured using size-exclusion chromatography (Fig. 5C-F, Table S5A). Both methods/growth conditions showed that the ss1-16 and ss1-17 mutants accumulated ∼3% less amylopectin (and consequently, 3% more amylose) than their wild-type controls.

**Fig. 5.**
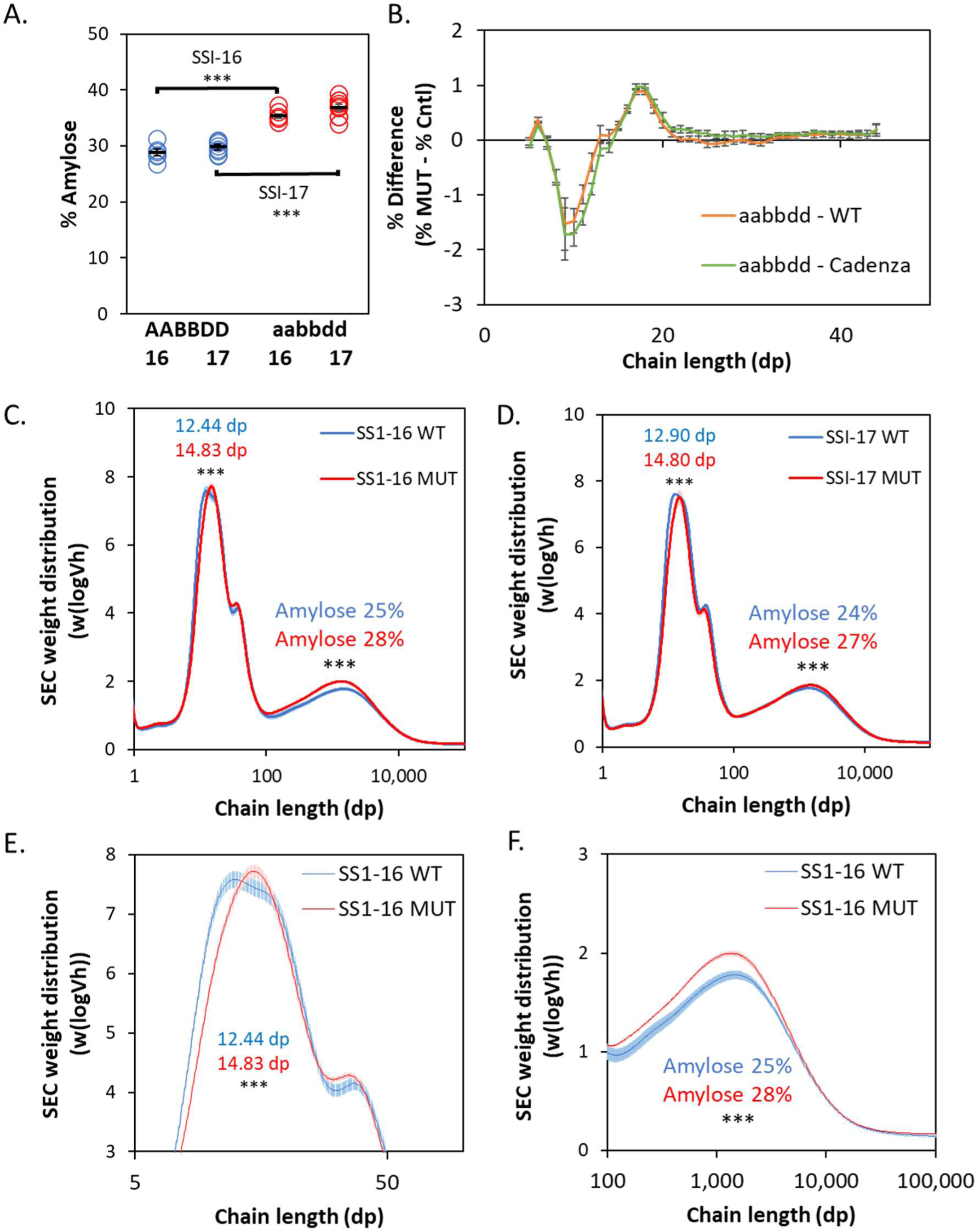
Starch molecular structure. Starches were purified from the grains of (A) glasshouse-grown or (B-F) field-grown plants (2023). The values are means ± SE of six replicate starches, each from the grains of different plants (A) or plots (B-F). Statistical differences between mutant and wild type were determined using Student’s t-tests. Wild type (WT/AABBDD) are in blue. Mutant (MUT/aabbdd) are in red. **A.** Amylose content was measured using an iodine-binding assay. **B.** Amylopectin chain length distributions of debranched starches were measured using anion-exchange chromatography. For each chain length, the difference (as % total starch) between the ss1-16 mutant and wild-type control (orange) or between the ss1-16 mutant and Cadenza parent (green) was calculated. Values are means. Negative values indicate a deficiency in the mutant. **C, D** Chain-length distributions of debranched starches were measured by size-exclusion chromatography (SEC). Data are for SS1-16 (C) and SS1-17 (D). For each starch, amylose content was estimated as the proportion of chains >100 DP. The values for average (mode) chain length (DP) for the amylopectin peak are indicated. **E, F** Data are as in (C) but the amylopectin (E) and amylose (F) peaks are expanded for clarity.

The amylopectin structure of purified and debranched starches from the 2023 field plots was determined using anion exchange chromatography (Dionex, Fig. 5B) and size-exclusion chromatography (SEC, Fig. 5C-F). Both methods revealed a relative deficiency in the shortest amylopectin chains in the ss1 mutants. Anion exchange chromatography (which gives a higher resolution of chain lengths than SEC) showed an increase in the abundance of chains of 6-7 DP and a reduction in chains of 8-12 DP. This suggests that the substrates for SS1 are 6-7 DP and that SS1 elongates these chains to 8-12 DP. The SEC results show that, due to reduced amounts of short chains in the ss1 mutant starches, the average amylopectin chain length was increased from 13 to 15 DP.

### ss1 mutant starch and flour have unique functional properties

To compare the gelatinization of mutant and wild-type starches, we observed purified starches in dilute Lugol’s solution as they were heated from 25-90 °C using a microscope with a heated stage, λ-plate and polarizing filters (Fig. 6A). By 65 °C, both types of starch had begun to lose birefringence, and this was entirely lost by 75 °C. The mutant starch had less birefringence than the wild type at 70 °C. This suggests that ss1 mutant starch is less thermally stable.

**Fig. 6.**
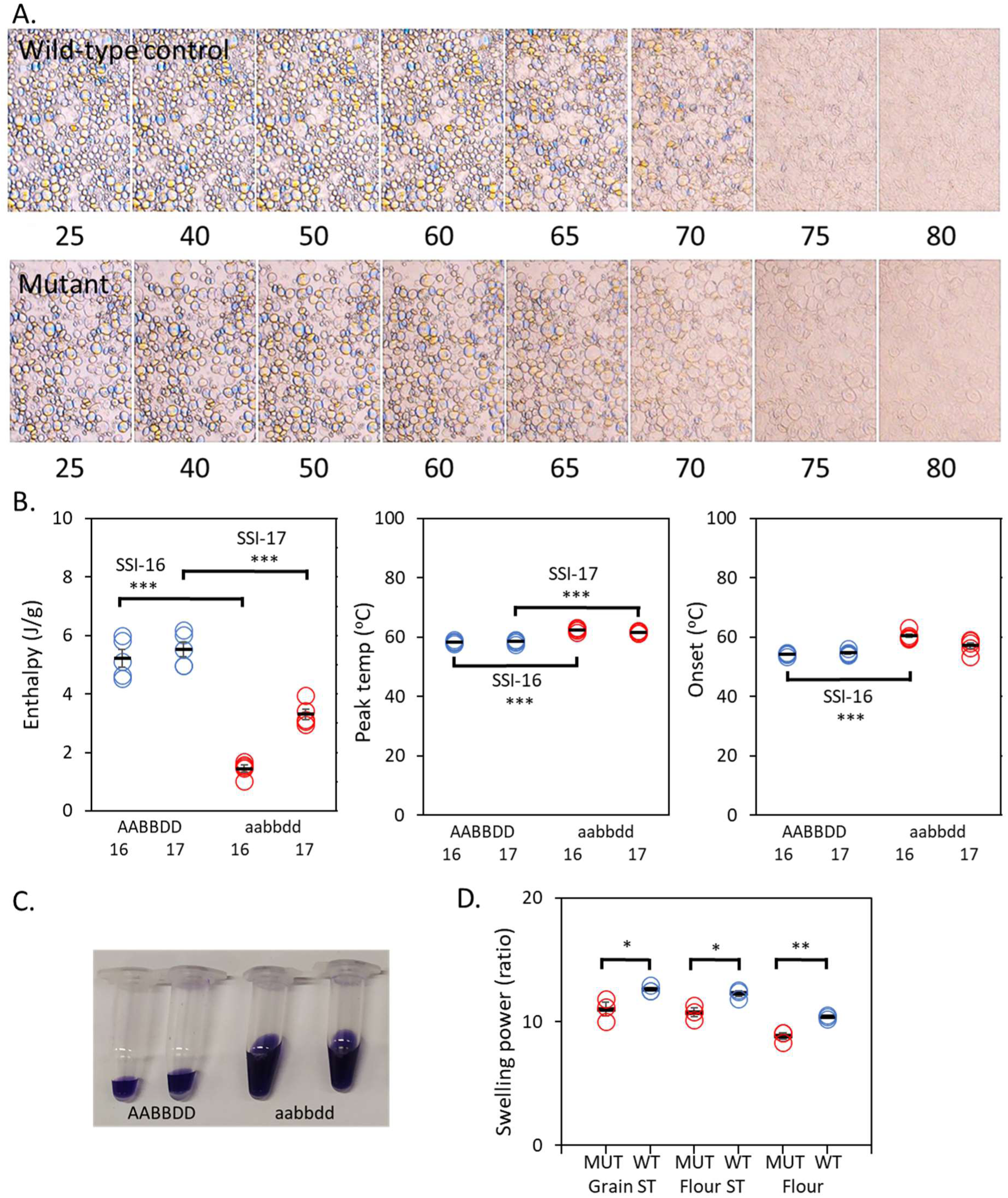
Starch functional properties. Starches were purified from the grains of field-grown plants. **A.** Starches purified from SS1-17 mutant or wild-type control white flours were suspended in dilute Lugol’s solution on a glass slide and heated at 5 °C per min using a microscope with heated stage. Images were taken at intervals using a tint plate and polarizing filters to observe loss of birefringence with temperature. **B.** Thermal properties were measured using differential scanning calorimetry (DSC). Values are means ± SE of five replicate starches, each from grains from a different field plot, together with the individual data for each starch. Statistical differences between mutant and wild type were determined using Student’s t-tests. Differences between mutant and corresponding wild-type controls that were not significant are not shown. Wild types (AABBDD) are in blue. Mutants (aabbdd) are in red. **C.** Samples for SDS-PAGE (Fig. S1C) were prepared by gelatinizing starches in gel sample buffer containing 2 % (w/v) LDS. The pellets after centrifugation are shown on the left. The ss1 mutant (aabbdd) samples swelled noticeably more the those of the wild type (AABBDD). **D.** Starch swelling power in water was measured using white flour, and two different starch preparations. Samples of wild type and mutant field-grown SS1-17 grain (2024) were milled to white flour. Starch was extracted from some of this white flour. Starch was also extracted from the original grain samples.

The thermal properties of SS1 wild-type and mutant starches were quantified using differential scanning calorimetry (Fig. 6B). There were statistically significant differences between wild-type and mutant starches of both ss1-16 and ss1-17 for enthalpy and peak temperature, whilst the onset temperature was significantly different for ss1-16 starches only. Enthalpy was reduced in the mutants whereas the peak and onset temperatures were increased.

During preparation of starch granule-bound proteins for SDS-PAGE (Fig. S1D), a noticeable difference in the volume of the gelatinized starches was observed between ss1 mutant and wild-type starches (Fig. 6C). This suggested that starch swelling power was affected by the ss1 mutations. The swelling power of SS1-17 flour and starch was quantified by gelatinization in water, rather than in SDS, as water is a more functionally relevant solute (Fig. 6D). In contrast to the higher swelling power of the mutant starches in SDS, in water, the swelling power of ss1-17 mutant flour, and starches purified from grains or from white flour, was lower than that of the wild-type controls.

Increases in amylose content can influence the rate and extent of amylolysis. To determine whether the small increase in amylose content in ss1 mutants impacted on amylolysis, we measured the digestibility of wholemeal flour in vitro using pancreatic α-amylase (Supplementary Fig. S4). The assay conditions used were those that have been shown to predict the human glycaemic response to the ingestion of starch-rich cooked food (Edwards *et al*., 2019). However, our results did not reveal measurable differences between ss1-16 mutants and wild-type controls in the digestibility of raw, cooked, or cooked and cooled wholemeal flours.

## Discussion

Our results show that the absence of SS1 in wheat leads to mild but significant changes in starch molecular structure. In agreement with previous studies of SS1-deficient plants (Delvallé et al., 2005; Fujita et al., 2006; McMaugh et al., 2014), we found a specific reduction in the proportion of the shortest amylopectin chains (corresponding to A1 chains). This confirms that SS1 has a unique role in synthesizing chains of this length, a function that other SS isoforms are unable to perform.

In their most severe mutant, McMaugh *et al*., (2014) found that RNAi suppression of ss1 in wheat resulted in a ∼10% reduction in grain starch content, and that this was due to decreased amylopectin synthesis. In our studies of ss1 Cadenza TILLING mutants, the amylopectin content of the mutants was reduced but the amylose content increased, such that the starch content of ss1 grains was near normal. This suggests that in our ss1 mutants, increased amylose synthesis can largely compensate for reduced amylopectin synthesis. A similar, but opposite, compensatory mechanism has been observed in Granule-Bound Starch Synthase (GBSS) mutants of wheat that entirely lack amylose: they also have near-normal starch contents showing that amylopectin synthesis can compensate for reduced amylose synthesis (Zang et al., 2012). In contrast, wheat TILLING mutants lacking other isoforms of starch synthase have both reduced grain weight and starch content (SS2a; Botticella et al., 2016; Schoen et al., 2021 and SS3a, Fahy et al., 2022).

The subtle effects of SS1-deficiency on starch content and grain weight in wheat and rice may explain why an SS1-deficient mutant has not, so far, been described for maize, even though SS1 accounts for approximately 60% of the total soluble SS in developing maize endosperm (Cao *et al*., 1999). Most classic maize starch mutants (such as dull1, which lacks Starch Synthase 3) (Cao et al., 1999) were found in forward genetic screens, because they gave a visual phenotype (i.e. shrunken grains due to low starch content). In contrast, the rice and wheat ss1 mutants were both obtained in reverse genetic screens, and both have little or no impact on plant, or grain, growth and development. The rice mutant also has starch granules that are normal in shape and crystallinity (Fujita *et al*., 2006). Given the relatively modest phenotypic defects of ss1 cereal mutants, and the existence of a potentially deleterious mutation in SS1-7A in an elite cultivar of wheat, leads us to question the adaptive significance of SS1 in plants. SS1 is not unique amongst starch synthases in this respect: the same lack of known adaptive significance also applies to the enzyme responsible for amylose synthesis, GBSS (Seung, 2020).

Increased amylose content in cereal grains is valued because of the associated increase in resistant starch, an important component of dietary fibre. Flour and bread produced from mutants lacking Starch Branching Enzyme 2 (SBE2), SS2a, or SS3a have amylose contents that are greater than controls by 4.5% (ss2a; Schoen et al., 2021) to 44.7 % (sbe2b, Botticella et al., 2018), and they have shown significantly higher resistant starch content and/or lower starch digestibility (Botticella et al., 2018; Corrado et al., 2022; Fahy et al., 2022; Schoen et al., 2021). SS1-deficient mutants also have increased amylose content (from 29.4 to 36.1 % i.e. 6.8% greater than control) but when tested, this had no effect on the digestibility of starch in wholemeal flour (either raw or hydrothermally processed). Thus, despite the increase in amylose content in ss1 mutant wheat flour being within the range seen for other starch mutants, the starch was digested at a normal rate in vitro. We suggest that abnormalities in amylopectin structure specific to the sbe2, ss2a and ss3a wheat mutants, may contribute to reduced starch digestibility.

Grain composition predicted by NIR analysis of whole grains of the ss1 mutants compared to wildtype controls showed no consistent differences between lines for starch and protein contents, and no significant differences for NDF (neutral detergent fibre) contents. Grain shape in the mutants was also normal. The normal whole grain NDF contents predicted by NIR analysis, of ss1 mutant grains contrasts with the biochemical measurements for dietary fibre components (total arabinoxylan (AX) and mixed-linked β-glucan (MLG)) seen in wholemeal flour from other wheat starch mutants (GBSS, SBE2 and SS2a: Botticella et al., 2018; SS3a: Fahy et al., 2022), where lower grain starch content and smaller grains correlated with increased fibre contents. The increased proportion of outer bran layers to endosperm in these small-grained mutants may explain the increase in these fibre components.

Biochemical analysis of white flour showed that the ss1 mutants had ∼20% more of the major dietary fibre components of wheat endosperm, AX and MLG than wildtype controls. White flour does not contain the outer bran layers (unlike wholemeal flour) and the AX and MLG contents reflect the cell wall composition of the endosperm. Higher fibre levels in white flour are desirable as they are associated with a wide range of health-promoting effects (Prins and Kosik, 2023) and they may also enhance food processing and product quality (Rosicka-Kaczmarek et al., 2016). Although the AX content of both ss1 mutant and wild type white flours were within the values reported for a range of wheat varieties (Ordaz-Ortiz, Devaux and Saulnier, 2005; Gebruers et al., 2010), understanding the mechanism behind the increase in fibre in the endosperm of ss1 mutants might aid future attempts to breed higher-fibre wheats without a significant yield penalty.

An observation common to wheat starch mutants, whether they lead to low starch content or not, is that they have increased contents of sucrose and sometimes other soluble carbohydrates in the mature grains (Botticella et al., 2018). The ss3a mutant also had increased total sugar (sucrose, glucose and fructose) content in immature grains (25 DAA) (Fahy et al., 2022.). Since soluble carbohydrates are the substrates for both fibre synthesis as well as starch, competition for the same substrate pool may explain the increased fibre content in the mutant endosperm. Measurement of starch, sugar and fibre contents in developing endosperms of SS1 and other starch mutants would allow this theory to be tested. Alternatively, at present we cannot completely rule out the involvement of a genetic factor other than SS1 that differs between mutant and wild type in both lines and independently controls fibre content. We studied two SS1 lines to reduce the chances of additional mutant genes being common to both lines but the existence of co-inherited haplotype blocks in wheat places limits on this approach (Brinton et al., 2020).

Although the effects of ss1 mutations on plant growth are minor, the changes to the functional properties of starch and flour may be industrially significant. Small changes in starch granule morphology and the rheological properties of flour (due to modified starch molecular structure and/or fibre content) can have large effects on milling and baking performance. Here, we found that both mutant lines, ss1-16 and ss1-17, exhibited a slightly lower proportion of A-type granules (i.e. higher % B-type) compared to their respective wild types (Table S4). When gently extracted from the mature grains, the A-type starch granules in the mutants were predominantly normal in diameter although grinding to wholemeal, and particularly to white flour, prior to starch extraction increased the proportion of larger-than-normal granules (Fig. 4). Microscopy also showed that the grinding of grains to flour appeared to increase the number of unusually large A-type granules with reduced birefringence (Fig. 3). Similarly, McMaugh et al. (2014) saw some deformed A-granules in their most severe SS1 RNAi wheat line. Together, these results suggest that, compared to wild-type controls, the A-type starch granules in ss1 mutant wheat grains may be more prone to physical damage during milling resulting in deformed A-granules with reduced crystallinity.

Despite their largely normal granule morphology, the thermal properties of ss1 mutant and wild type starches were distinctly different, suggesting that these are predominantly determined by molecular structure. When gelatinized in water, starch swelling power in the ss1 mutants was reduced and there were significant differences between wild-type and mutant starches in DSC gelatinization parameters (enthalpy, onset temperature and peak temperature). In both mutant lines, enthalpy was reduced, whilst onset and peak temperatures were increased. Enthalpy is the amount of heat needed to gelatinize starch granules (and it reflects the level of molecular organization of the starch), whilst onset and peak temperatures indicate the temperatures needed to initiate and achieve gelatinization, respectively. Reduced enthalpy suggests that the starch in the ss1 mutants may be less crystalline (more amorphous). However, increased peak and onset temperatures suggest that the starch crystals have increased thermal stability, possibly due to increased structural order. Our swelling power and DSC results broadly agree with those of McMaugh et al. (2014). They found that suppression of *SS1* expression altered starch gelatinization properties, including reduced swelling power, and higher peak and end gelatinization temperatures and lower enthalpy.

Further studies of ss1 mutant wheat, including assessments of field performance and end-use functionality (as determined by both starch and fibre), will be essential to determine whether ss1 mutants offer advantages for commercial applications. Starch gelatinization, which occurs during the thermal processing of starchy foods, plays a critical role in determining the quality of food products. Cell wall polysaccharides also significantly contribute to flour performance by influencing water absorption, dough rheology, and baking parameters. If they are shown to enhance food processing and product quality, the combination in ss1 mutant wheat of alterations in functional characteristics with near-normal grain yield could be very interesting.

## Supporting information

Supplemental informatiom

## Supplementary data

The following supplementary data are available at JXB online.

**Table S1.** SS1 null mutations in the hexaploid wheat variety, Cadenza identified in-silico.

**Table S2.** KASP primers used for marker-assisted selection of SS1 mutant and wild-type plants.

**Table S3.** Compositional and morphometric characteristics of field-grown grains measured by NIR and Marvin grain analyser.

**Table S4.** Starch granule-size distribution of purified starches.

**Table S5.** Chain-length distribution determined by SEC of purified starches.

**Figure S1.** Diagrams of *SS1* gene structure and the crossing scheme used to develop mutant lines.

**Figure S2.** *SS1* expression pattens according to the exp-VIP database.

**Figure S3.** SDS-PAGE gels of proteins from purified starches and Western blots of starch granule-bound and soluble proteins.

**Figure S4.** The in vitro digestibility of SS1 wild type and mutant wholemeal flour.

## Acknowledgements

We thank the following people and groups for their contributions to this publication: the John Innes and Quadram Institutes Bioimaging facilities and staff (particularly Rhea Stringer, Eva Wegel, James Lazenby and Catherine Booth) for imaging and training, the JIC Horticultural Services team for raising our glasshouse plants, the JIC Field Experimentation Team for carrying out our field trials, Martin Radjzek and Baldeep Kular are thanked for Dionex training and Robert Hindmarsh is thanked for training and access to DSC equipment at the University of East Anglia, Norwich, UK.

## Authors contribution

KT, and BAH*: conceptualization;* KT, and BAH*: methodology;* KT, BF, OG, MP, and OK*: formal analysis;* KT, BF, OG, MP, JHA-H, JM, and OK*: investigation;* KT*: data curation;* KT, and BAH*: writing - original draft;* KT, BF, OG, MP, JHA-H, JM, OK, AL, FW, and BAH*: writing - review & editing;* KT, BF, MP, and OK*: visualization;* AL, FW, and BAH*: supervision;* BAH*: funding acquisition*.

## Conflicts of interest

No conflict of interest declared.

## Funding

This work was supported by the Biotechnology and Biological Sciences Research Council (BBSRC) under the following Institute Strategic Programmes (ISPs): Food Innovation and Health [grant number BB/R012512/1 and its constituent projects BBS/E/F/000PR10343 and BBS/E/F/000PR10345]; Molecules from Nature – Crop Quality [grant number BBS/E/J/000PR9799]; Designing Future Wheat [grant number BB/P016855/1], Delivering Sustainable Wheat [grant number BB/X011003/1].

## Data availability

The datasets generated and analysed during the current study (data for Trafford et al. 2026; DOI: 10.5281/zenodo.18682877) are available in the Zenodo DSW Data Repository at https://zenodo.org/communities/dsw.

## Abbreviations

DP: Degree of polymerization
DSC: Differential scanning calorimetry
HPLC: High performance liquid chromatography
FWT: Fresh weight
DWT: Dry weight
SEC: Size exclusion chromatography
SEM: Scanning electron microscopy
TILLING: Targeting induced local lesions in genomes
AX: Arabinoxylan
TO-AX: Total AX
WE-AX: Water-extractable AX
AXOS AX: oligosaccharides
MLG: Mixed linkage glucan (β-glucan)
MLGOS: MLG oligosaccharides
WT: Wild type (AABBDD)
MUT: Mutant (aabbdd)
NIR: Near-infrared

## Notes

https://zenodo.org/communities/dsw

### Competing Interest Statement

The authors have declared no competing interest.

