## Supplemental informatiom for "Wheat mutants lacking Starch Synthase 1 have altered starch composition and cell wall content"

### Supplementary data

**Table S1. *SS1* alleles.**

| Gene | Sub genome | Line | Variant ID | Consequence | Alleles | cDNA coordinate | Amino acids | Amino acid coordinate |
|---|---|---|---|---|---|---|---|---|
| TraesCS7A02G120300 | A | 16+17 | C0451 | stop | G/A | cDNA 1525 | R/* | 396 |
| TraesCS7B02G018600 | B | 16 | C0169 | stop | G/A | cDNA 1180 | W/* | 159 |
| TraesCS7B02G018600 | B | 17 | C1533 | stop | G/A | cDNA 2239 | W/* | 512 |
| TraesCS7D02G117800 | D | 16+17 | C2085 | stop | G/A | cDNA 1938 | W/* | 646 |

Information for A and B homoeologs is from Ensembl Plants release 60 - October 2024 © EMBL-EBI. Information on Cadenza2085, and all four genotypes are available from www.seedstor.ac.uk.

**Table S2. KASP primers.**

| Gene | Sub genome | Line | Variant ID | VIC-wild type | FAM-mutant | Common |
|---|---|---|---|---|---|---|
| TraesCS7A02G120300 | A | 16+17 | C0451 | GAAGGTCGGAGTCAACGGATTcctcaatgagctcttaagttccC | GAAGGTGACCAAGTTCATGCTcctcaatgagctcttaagttccT | acccaaagagcaaactgaattaatt |
| TraesCS7B02G018600 | B | 16 | C0169 | GAAGGTCGGAGTCAACGGATTgcatgtttgttgtgaacgattgG | GAAGGTGACCAAGTTCATGCTgcatgtttgttgtgaacgattgA | actttcagatccacaaacaacg |
| TraesCS7B02G018600 | B | 17 | C1533 | GAAGGTCGGAGTCAACGGATTacgagcagatcttcgagtgG | GAAGGTGACCAAGTTCATGCTacgagcagatcttcgagtgA | ctcaaggagactcgcacttg |
| TraesCS7D02G117800 | D | 16+17 | C2085 | GAAGGTCGGAGTCAACGGATTgtacgagcagatcttcgaatgG | GAAGGTGACCAAGTTCATGCTgtacgagcagatcttcgaatgA | tcttcagagctcaaggagacc |

Sequences are 5’ to 3’ VIC/FAM tail sequences (capital letters) are at the 5’ end and the SNP is at the 3’end of the primer sequence.

**Table S3. Grain characteristics.**

**A.**

| Line | Method | Phenotype | Genotype | Mean | SE | Student's t-test (p-value) |
|---|---|---|---|---|---|---|
| SSI-16 | NIR | Predicted Hardness (empirical) | WT | 76.33 | 3.54 | ** |
| SSI-16 | NIR | Predicted Hardness (empirical) | Mutant | 62.11 | 2.00 | |
| SSI-16 | NIR | Predicted Moisture (%) | WT | 13.99 | 0.22 | |
| SSI-16 | NIR | Predicted Moisture (%) | Mutant | 14.31 | 0.17 | |
| SSI-16 | NIR | Predicted NDF (Dry basis, %) | WT | 16.43 | 0.54 | |
| SSI-16 | NIR | Predicted NDF (Dry basis, %) | Mutant | 16.62 | 0.93 | |
| SSI-16 | NIR | Predicted Protein (Dry basis, %) | WT | 16.50 | 0.36 | ** |
| SSI-16 | NIR | Predicted Protein (Dry basis, %) | Mutant | 14.86 | 0.32 | |
| SSI-16 | NIR | Predicted Starch (Dry basis, %) | WT | 66.79 | 0.40 | |
| SSI-16 | NIR | Predicted Starch (Dry basis, %) | Mutant | 67.70 | 0.51 | |
| SSI-16 | Marvin | Grain weight (mg) | WT | 52.55 | 1.30 | * |
| SSI-16 | Marvin | Grain weight (mg) | Mutant | 47.97 | 1.38 | |
| SSI-16 | Marvin | Area ($mm^2$) | WT | 21.05 | 0.42 | |
| SSI-16 | Marvin | Area ($mm^2$) | Mutant | 20.87 | 0.35 | |
| SSI-16 | Marvin | Width (mm) | WT | 3.72 | 0.04 | |
| SSI-16 | Marvin | Width (mm) | Mutant | 3.60 | 0.03 | |
| SSI-16 | Marvin | Length (mm) | WT | 7.05 | 0.08 | |
| SSI-16 | Marvin | Length (mm) | Mutant | 7.13 | 0.08 | |
| SSI-17 | NIR | Predicted Hardness (empirical) | WT | 74.17 | 0.84 | *** |
| SSI-17 | NIR | Predicted Hardness (empirical) | Mutant | 66.83 | 1.17 | |
| SSI-17 | NIR | Predicted Moisture (%) | WT | 13.83 | 0.11 | * |
| SSI-17 | NIR | Predicted Moisture (%) | Mutant | 14.29 | 0.15 | |
| SSI-17 | NIR | Predicted NDF (Dry basis, %) | WT | 15.98 | 0.78 | |
| SSI-17 | NIR | Predicted NDF (Dry basis, %) | Mutant | 15.33 | 0.65 | |
| SSI-17 | NIR | Predicted Protein (Dry basis, %) | WT | 14.63 | 0.44 | |
| SSI-17 | NIR | Predicted Protein (Dry basis, %) | Mutant | 15.45 | 0.18 | |
| SSI-17 | NIR | Predicted Starch (Dry basis, %) | WT | 69.23 | 0.50 | * |
| SSI-17 | NIR | Predicted Starch (Dry basis, %) | Mutant | 67.81 | 0.39 | |
| SSI-17 | Marvin | Grain weight (mg) | WT | 39.69 | 2.30 | |
| SSI-17 | Marvin | Grain weight (mg) | Mutant | 44.18 | 1.27 | |
| SSI-17 | Marvin | Area ($mm^2$) | WT | 15.85 | 0.48 | |
| SSI-17 | Marvin | Area ($mm^2$) | Mutant | 16.98 | 0.37 | |
| SSI-17 | Marvin | Width (mm) | WT | 3.28 | 0.09 | |
| SSI-17 | Marvin | Width (mm) | Mutant | 3.44 | 0.05 | |
| SSI-17 | Marvin | Length (mm) | WT | 6.58 | 0.03 | |
| SSI-17 | Marvin | Length (mm) | Mutant | 6.66 | 0.06 | |

All plants were grown at the same time in 2023 in randomized plots. Values are means of six biological replicates, each from a different field plot. For NIR, each grain sample was analysed three times. NDF = neutral detergent fibre (a prediction of the insoluble plant cell wall content). Statistical differences between mutant (aabbdd) and wild type (WT, AABBDD) were determined using Student’s t-tests.

**B. Calibration parameters for the Perten DA 7250 NIR Analyser.**

The data below were provided by the manufacturer in application Note DA Wheat.

| **Parameter** | **Range (%)** | **Samples** | **R** |
|---|---|---|---|
| Hardness (SKCS) (HI) | 12.9-87.2 | 100+ | 0.81 |
| Moisture | 7.3-22.1 | 4000+ | 0.97 |
| NDF | 7.2-17.4 | <100 | 0.77 |
| Protein | 8.2-22.8 | 4200+ | 0.98 |
| Starch | 61.5-83.0 | 1200+ | 0.97 |

**Table S4. Starch granule-size distribution.**

| Line | Genotype | A-granule diameter (µm) | SE | Student's t-test | B-granule diameter (µm) | SE | Student's t-test | B granule content (%) | SE | Student’s t-test |
|---|---|---|---|---|---|---|---|---|---|---|
| SS1-17 | AABBDD | 17.92 | 0.26 | 0.215 | 5.98 | 0.06 | 0.000 | 50% | 000 | 0040 |
| SS1-17 | aabbdd | 18.30 | 0.12 | | 6.80 | 0.14 | *** | 83% | 001 | * |
| SS1-16 | AABBDD | 18.83 | 0.42 | 0.096 | 6.85 | 0.25 | 0.229 | 123% | 001 | 0004 |
| SS1-16 | aabbdd | 17.99 | 0.19 | | 7.29 | 0.24 | | 186% | 001 | ** |

All plants were grown at the same time in 2023 in randomized field plots. Purified starches were subjected to analysis of granule-size distribution using a Multisizer 4e Coulter counter. Values are means of six biological replicates, each from a different field plot. Statistical differences between mutant (aabbdd) and wild type (AABBDD) are indicated. Only the B-granule content was consistently different for both lines, being higher in the ss1 mutant.

**Table S5. Chain-length distribution determined by SEC.**

| Line | Genotype | Amylopectin chain length (peak DP) | SE | Student's t-test | Amylose content (% total starch) | SE | Student's t-test |
|---|---|---|---|---|---|---|---|
| SS1-16 | AABBDD | 12.44 | 0.05 | 0.000 | 24.89% | 0.34% | 0.000 |
| SS1-16 | aabbdd | 14.83 | 0.04 | *** | 27.57% | 0.17% | *** |
| SS1-17 | AABBDD | 12.90 | 0.05 | 0.000 | 24.36% | 0.17% | 0.000 |
| SS1-17 | aabbdd | 14.80 | 0.10 | *** | 26.60% | 0.08% | *** |

All plants were grown at the same time in 2023 in randomized field plots. Purified starches were subjected to analysis of chain-length distribution using size-exclusion chromatography. From these data, the peak amylopection chain length (DP) and the amylose content as a percentage of total starch were calculated. Values are means of six biological replicates, each from a different field plot. Statistical differences between mutant (aabbdd) and wild type (AABBDD) are indicated.

**Figure S1. Selection of ss1 triple null mutants.**

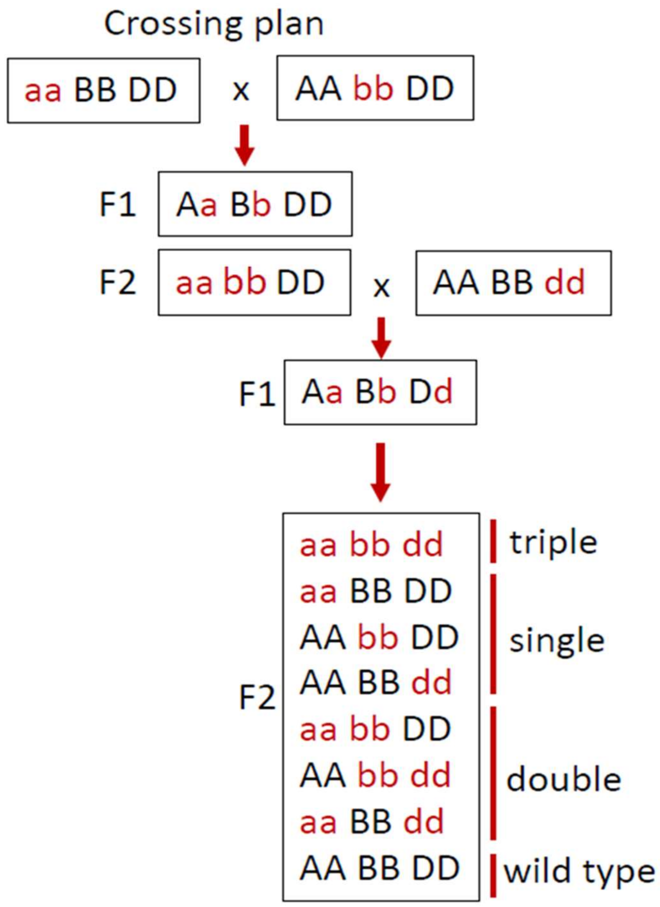


To generate ss1 mutant lines 16 and 17 (which have different *SS1-7B* mutations, Table S1), the A-genome mutant was crossed to each B-genome mutant, AB double mutants were selected and then they were each crossed with the D-genome mutant. Triple heterozygous $F_1$ plants were identified (from two crosses, 16 and 17), and the various genotypes of interest were identified from the $F_2$ progenies.

**Figure S2. Expression analysis.**

Expression data from the expVIP database for the three homoeologues genes of SS1 are shown (https://www.wheat-expression.com). Values are tpm ± SE for the number of studies indicated except for studies with only one replicate value, where no SE values is given (na = not applicable). Colour scales have been applied to the tpm values in each table separately.

**A.** Comparison between tissue types.

**B.** Data from different studies for expression in endosperm, starchy endosperm and aleurone plus starchy endosperm.

**C.** Data for endosperm at different stages of development.

**A.**

| **Tissue type** | **TraesCS7A02G120300** | | **TraesCS7B02G018600** | | **TraesCS7D02G117800** | |
|---|---|---|---|---|---|---|
| **Gr>Lf>Sp>Ro** | **tpm** | **SE** | **tpm** | **SE** | **tpm** | **SE** |
| Roots (n=89) | 1.14 | 1.20 | 1.41 | 2.00 | 2.03 | 1.68 |
| Spike (n=280) | 12.16 | 7.11 | 16.75 | 10.22 | 20.69 | 12.76 |
| Leaves/shoots (n=481) | 16.01 | 11.38 | 29.34 | 19.95 | 20.05 | 12.66 |
| Grain (n=166) | 18.50 | 12.03 | 40.26 | 24.33 | 28.32 | 21.70 |

**B.**

| **Endosperm** | **TraesCS7A02G120300** | | **TraesCS7B02G018600** | | **TraesCS7D02G117800** | |
|---|---|---|---|---|---|---|
| **7B>7D>7A** | **tpm** | **SE** | **tpm** | **SE** | **tpm** | **SE** |
| Endosperm (n=3) | 18.34 | 2.20 | 37.72 | 3.90 | 24.25 | 4.69 |
| Endosperm (n=4) | 25.20 | 11.27 | 43.18 | 22.87 | 28.48 | 17.54 |
| Endosperm (n=14) | 30.72 | 21.55 | 54.03 | 40.56 | 44.96 | 44.58 |
| Endosperm (n=8) | 35.60 | 16.45 | 61.37 | 25.41 | 46.87 | 25.41 |

**C.**

| **Endosperm (age, dpa)** | **TraesCS7A02G120300** | | **TraesCS7B02G018600** | | **TraesCS7D02G117800** | |
|---|---|---|---|---|---|---|
| **6-14 dap>20-30 dpa** | **tpm** | **SE** | **tpm** | **SE** | **tpm** | **SE** |
| 6 dpa (n=1) | 55.59 | na | 55.95 | na | 76.52 | na |
| 9 dpa (n=1) | 59.29 | na | 103.24 | na | 107.79 | na |
| 10 dpa (n=4) | 47.52 | 7.11 | 78.05 | 15.65 | 66.79 | 11.42 |
| 12 dpa (n=3) | 15.28 | 0.75 | 31.38 | 1.62 | 24.29 | 0.34 |
| 14 dpa (n=1) | 82.83 | na | 172.11 | na | 167.27 | na |
| 20 dpa (n=4) | 32.72 | 2.18 | 57.69 | 4.21 | 39.97 | 1.86 |
| 20 dpa (n=4) | 23.67 | 14.21 | 44.69 | 22.79 | 26.95 | 17.83 |
| 30 dpa (n=4) | 13.90 | 1.72 | 25.08 | 2.27 | 11.29 | 1.60 |
| 30 dpa (n=4) | 25.20 | 11.27 | 43.18 | 22.87 | 28.48 | 17.54 |

**Figure S3. Analysis of SS1 distribution between soluble and starch granule-bound fractions.**

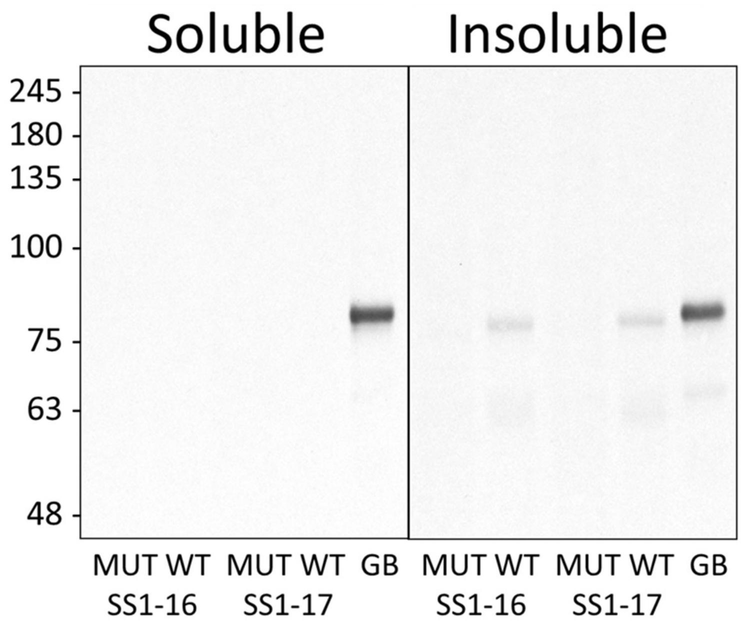


Developing grains of SS1-16 and SS1-17 mutants (MUT) and their wild-type controls (WT) were extracted and divided into soluble and insoluble fractions with equivalent volumes. For each extract, equal proportions of soluble and insoluble proteins were separated using SDS-PAGE, blotted onto membranes, which were then probed with an SS1-specific antibody. A track of starch granule-bound proteins (GB) extracted from starch isolated from mature WT grains was included as a positive control. SS1 protein was detected in the insoluble fractions of wild type grains but not mutant grains. No SS1 was detected in the soluble fractions from equal proportions of the grain suggesting that, at this stage of development, SS1 is predominantly granule bound.

**Figure S4. Amylase digestibility of flour *in vitro.***

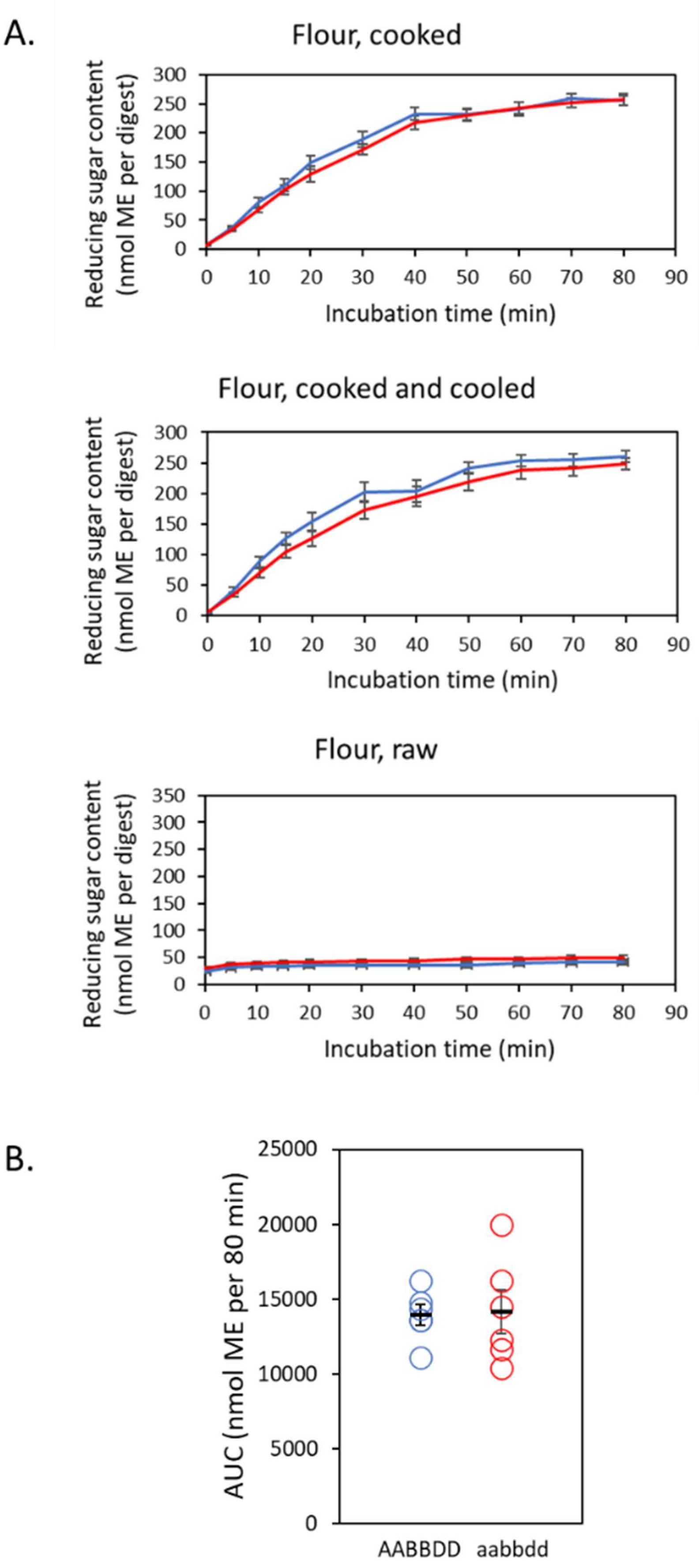

Wholemeal flour samples were prepared from wild type and mutant SS1-16 grains (2023). Replicate samples of flour in PBS were temperature-treated (cooked, cooked and cooled or raw), then subsampled prior to digestion with pancreatic α-amylase. The amylase reaction was stopped after 0-80 min and the reducing sugar content of the soluble fraction was determined. Reducing sugar content is expressed as maltose equivalents (ME). The values are means ± SE of six replicate flours, each from the grains of a different field plot. Each digest was duplicated. Wild types are in blue. Mutants are in red.

**A.** The reducing sugars released by amylase digestion of starch are shown as a function of time.

**B.** For each replicate cooked flour in (A), the area under the digest curve (AUC) was determined. In addition to the means ± SE, the individual values for each flour are shown. No statistical differences in digestibility between mutant and wild type were found using Student’s t-tests.
